# Integrated microbiome and metabolomic profiling reveals alterations across the adenoma–colorectal cancer sequence

**DOI:** 10.64898/2026.03.20.713264

**Authors:** Tien-En Chang, Hung-Hsin Lin, Jiing-Chyuan Luo, Ying-Fan Chen, Yen-Po Wang, Kuei-Chuan Lee, Pei-Chang Lee, Yi-Tsung Lin, Hui-Chun Huang, Chien-Wei Su, Yi-Hsiang Huang, Ming-Chih Hou

**Affiliations:** Endoscopy Center for Diagnosis and Treatment; Division of Gastroenterology and Hepatology; Division of Infectious Diseases; Division of General Medicine, Department of Medicine; Division of Colorectal Surgery, Department of Surgery; Department of Medical Research, Taipei Veterans General Hospital, Taipei, Taiwan; School of Medicine; Institute of Biomedical Informatics, National Yang Ming Chiao Tung University, Taipei, Taiwan

**Keywords:** Gut microbiota, 16s rRNA full-length sequencing, Functional prediction, Metabolomic, Colorectal cancer, Colon adenoma

## Abstract

The incidence of colorectal cancer (CRC) has been increasing in Taiwan and is associated with multiple risk factors, including aging, obesity, and dietary habits. Increasing evidence suggests that gut microbiota dysbiosis contributes to CRC development. This study aimed to characterize microbial and metabolic alterations across premalignant and malignant colorectal lesions and to identify potential microbiome-associated biomarkers. Individuals undergoing colonoscopy for screening or surveillance at Taipei Veterans General Hospital were enrolled. Gut microbial composition was analyzed using full-length 16S rRNA gene sequencing to achieve high-resolution taxonomic profiling. Predicted functional pathways were inferred from microbial communities, and targeted metabolomic profiling was performed to evaluate microbial metabolic outputs.

A total of 122 individuals were included, comprising 62 healthy controls, 15 adenoma cases, and 45 CRC cases. Progressive shifts in microbial composition and predicted functional pathways were observed along the adenoma–carcinoma sequence. Several bacterial taxa, including *Phocaeicola dorei*, *Anaerotignum faecicola*, *Negativibacillus massiliensis*, and *Dysosmobacter segnis*, were enriched in CRC. At the functional level, CRC samples showed enrichment of pathways associated with energy metabolism and bacterial stress responses. Metabolomic analysis further revealed increased levels of tauro-ursocholanic acid in CRC samples, whereas short-chain fatty acids (SCFAs) were reduced compared with controls.

Integrative analysis combining full-length 16S sequencing, functional pathway prediction, and metabolomic profiling revealed coordinated microbial and metabolic alterations across the adenoma–carcinoma sequence. These findings provide insight into microbiome-associated processes in colorectal tumorigenesis and suggest potential microbial and metabolic biomarkers for CRC.

**Importance:** Colorectal cancer (CRC) develops through a adenoma–carcinoma sequence, yet the microbial and metabolic alterations accompanying this progression remain incompletely understood. In this study, we integrated full-length 16S rRNA gene sequencing with metabolomic profiling to characterize taxonomic, functional, and metabolic changes across healthy controls, adenoma, and CRC. Our results reveal synchronized shifts in specific microbial taxa, predicted metabolic pathways, and fecal metabolites along the adenoma–carcinoma sequence. Several bacterial species, including *Phocaeicola dorei*, *Anaerotignum faecicola*, and *Dysosmobacter segnis,* increased in CRC, whereas short-chain fatty acids decreased progressively from controls to adenoma and CRC. Functional pathway analysis further indicated alterations in microbial fermentation, amino acid metabolism, and energy-related pathways. Together, these findings highlight the potential role of microbiome-associated metabolic changes in colorectal tumorigenesis and suggest candidate microbial and metabolic markers that may aid in understanding disease development and improving risk stratification.

## Introduction

Colorectal cancer (CRC) has become the leading cancer in Taiwan in recent years. Screening for fecal occult blood, combined with colonoscopic detection and removal of precancerous lesions, has been demonstrated to significantly reduce the incidence and mortality associated with CRC. ^1^ The rising incidence of CRC can be attributed to factors such as aging, obesity, decreased physical activity, and dietary habits. ^2–4^ Beyond lifestyle factors, the development of CRC is closely linked to the accumulation of genetic alterations, a process commonly referred to as the adenoma-carcinoma sequence. ^5,6^ Gut microbiota and the microbial metabolites are believed to be involved in the process. ^2,3,5^

The gut microbiota maintain epithelial homeostasis, as commensal bacteria play a key role in promoting epithelial integrity. ^7^ Microbial dysbiosis is a well-documented phenomenon observed in patients with colonic adenoma and adenocarcinoma. For example, *Fusobacterium* is typically more abundant in CRC patients and patients with precancerous colorectal lesion compared to healthy individuals. ^5,8^ *Fusobacterium nucleatum* is found in significantly higher abundance in the stool samples of patients with adenomatous polyps compared to those from individuals with normal mucosa, hyperplastic polyps, or sessile serrated adenomas (SSA). ^8^ Similarly, *Proteobacteria*, which includes many pathogenic species, is also found in higher abundance in CRC patients. ^3^ A previous study demonstrated that *Bifidobacterium pseudocatenulatum* and *Faecalibacterium prausnitzii* exhibit a negative correlation with mutations in the adenomatous polyposis coli (APC) gene, whereas the abundance of *Fusobacterium mortiferum* is positively associated with APC mutations. ^9^ Thus, alterations in the gut microbiota may serve as valuable biomarkers for identifying patients with CRC or precancerous lesions, enabling earlier diagnosis and timely therapeutic intervention.

The fermentation of dietary components by anaerobic bacteria in the colon can generate a diverse collection of metabolites. ^2^ Short-chain fatty acids (SCFAs) are among the most widely discussed bacterial metabolites, produced through the fermentation of non-digestible carbohydrates, proteins and amino acids. ^2,10^ SCFAs contribute to gut barrier integrity by enhancing mucin expression and promoting the differentiation and apoptosis of colonic epithelial cells. SCFAs also have anti-inflammatory potential by inhibiting the activity of histone deacetylases. The presence and abundance of these metabolites reflect the functional capacity of the gut microbiota.

Currently, there is no consensus on the microbiota alterations associated with precancerous colon polyps and CRC compared to healthy individuals, as determined by full-length sequencing of the bacterial 16S rRNA gene and metagenomics in fecal samples. This study aimed to investigate alterations in the fecal microbiome and metabolomic profiles and to identify key microbial taxa or metabolites associated with premalignant and malignant colorectal lesions, where possible.

## Methods

### Patient selection

Subjects who underwent colonoscopy for screening or surveillance, including those with a positive fecal occult blood test, a history of colorectal polyps, or changes in bowel habits, were included in this study. After obtaining informed consent, stool samples were collected prior to bowel preparation. Anthropometric measurements were recorded, including height, weight, body mass index (BMI), waist circumference, and blood pressure. Blood samples were collected to assess metabolic syndrome parameters, including fasting glucose, total cholesterol, high-density lipoprotein (HDL), triglycerides, and glycated hemoglobin (HbA1c).

Subjects with severe cardiovascular, pulmonary, hepatic, or renal diseases, those with a history of gastrointestinal tract surgery, those with untreated underlying malignancies (except colorectal cancer), and vulnerable individuals—including aboriginal peoples, pregnant women, those with disabilities or mental illnesses, and residents of nursing homes or home care centers as well as those had taken proton pump inhibitors, nonsteroidal anti-inflammatory drugs, antibiotics, or probiotics within four weeks of sample collection were excluded. This study adhered to the standards of the Declaration of Helsinki and current ethical guidelines. The hospital’s Institutional Review Board (IRB) had approved the study (Approval No: 2021-12-003B).

After colonoscopic and histological examination, individuals with normal findings (Control_A), those with adenomas, and those with colorectal cancer (CRC) were included in the analysis. The CRC group was further subdivided into right-sided (CRC_R) and left-sided (CRC_L) tumors according to tumor location. Individuals presenting with multiple lesion types (e.g., adenoma, hyperplastic polyp, and CRC) were excluded; however, cases in which CRC was present were classified within the CRC group.

In addition, sequencing data from an independent cohort of healthy individuals were included as Control_B. These participants were screened fecal microbiota transplantation (FMT) donors who were confirmed to be healthy based on comprehensive blood and stool examinations, with no evidence of chronic illness, infectious disease, or gastrointestinal disorders. Although colonoscopic confirmation was not available, these individuals exhibited favorable metabolic characteristics, including lower body mass index and overall healthier metabolic profiles. ^11,12^

### Sample collection for metagenomic and metabolomics

In each subject, frozen fecal samples for metagenomic and metabolomics analysis were delivered to the hospital within four hours in insulated polystyrene foam containers before colonoscopy. All samples were stored at −80°C until further analysis to prevent bacterial overgrowth.

### Bacterial genomic DNA extraction and full-length 16S rRNA sequencing and analysis

Bacterial genomic DNA was extracted from fecal samples using the QIAamp Fast DNA Mini Kit (Qiagen, MD, USA) according to the manufacturer’s protocols. In brief, study samples weighing 180–220 mg yielded 5–100 μg of genomic DNA, which was directly used for 16S rRNA gene sequencing. The quantity and quality of the isolated genomic DNA were assessed using the NanoDrop ND-1000 (Thermo Scientific, Wilmington, DE, USA). Genomic DNA was stored at –80°C until 16S rRNA sequencing. One microliter of DNA (500 ng) was used as a template in a PCR reaction to amplify the full-length 16S rRNA gene (V1-V9 region) using barcoded forward and reverse primers. The barcoded 16S metagenomic samples were then pooled to create a single sample for SMRTbell library construction. The PCR procedure included an initial denaturation step at 95°C for 3 minutes, followed by 30 seconds at 95°C, annealing at 57°C for 30 seconds, and extension at 72°C for 1 minute. The final product was stored at 4°C. Next-generation sequencing was conducted using the PacBio Sequel System, SMRT Cell 1M, following the standard protocol. ^13–15^

### Targeted fecal metabolomic analysis for SCFA and bile acids (BAs)

Quantification of fecal SCFAs and BAs were determined in study subjects and the control by ultra-high performance liquid chromatography-mass spectrometry (UHPLC-MS/MS). ^16,17^ The samples were extracted from the collected stool. After suspension in double distilled water, homogenization, and centrifugation, the supernatant was taken for analysis.

The concentrations of SCFAs and BAs were compared between the groups. Linear regression was performed to evaluate the relationship between microbial metabolites, clinical metabolic parameters, and the alpha diversity indices.

To investigate the multivariate association structure between gut microbial genera and fecal metabolites, sparse partial least squares (sPLS) analysis via the mixOmics framework was performed. ^18,19^ The microbiome matrix consisted of genus-level CLR-transformed abundances of the ANCOM-BC2-selected taxa, and the metabolite matrix consisted of log-transformed and z-score standardized bile acid and short-chain fatty acid panel variables. Prior to integration, both matrices were residualized for group only, and the residualized matrices were used as inputs for sPLS. The model was fitted with two latent components and sparse feature selection was applied to both the microbial and metabolite blocks. Sample plots and variable plots were used to visualize the integrated structure and selected features.

### Data processing and statistical analysis

SMRT Link v10.1 was used to generate highly accurate reads (HiFi reads) with a read quality (RQ) >30, and to demultiplex the barcodes. DADA2 (version 1.20) was used for deduplication and chimera removal. After denoising, the amplicon sequence variants (ASVs) were processed using the QIIME2 v2023.9 pipeline. Alpha diversity was assessed through observed features index, Faith’s phylogenetic diversity (Faith PD), Pielou’s evenness index, and the Shannon diversity index (SI) within the QIIME2 framework. Taxonomic annotation was conducted using the Genome Taxonomy Database (GTDB, release v220). ^20^ Enterotype was calculated via online classifier Enterotyper, using the genus-level taxonomic profiles. ^21,22^

All statistical analyses of the bacterial community were performed using R software (v4.1.0). Constrained principal coordinates analysis (PCoA) on the Bray-Curtis distance and Aitchison distance matrix computed from centered log ratio transformation (CLR)-transformed data were performed. Permutational multivariate analysis of variance (PERMANOVA) were applied to compare between-group inertia percentages, and beta diversity was assessed using PERMDISP (betadisper with permutest), which evaluates whether distances of samples to their group centroids differ across groups. ^23^ The analysis of compositions of microbiomes with bias correction (ANCOM-BC2) was also applied to distinguish differentially abundant taxa between groups. ^24^ Welch’s t-test was performed to identify significant bacterial differences in pairwise comparisons.

Function prediction of microbial communities was performed using PICRUSt2 (version 2.4.1) based on 16S rRNA gene sequences. ^25^ ASVs were placed into a reference phylogenetic tree, and gene family abundances were inferred using KEGG orthologs (KOs) and Enzyme Commission (EC) numbers. ^26^

All data were expressed as means ± standard deviation. If some parameters were not normally distributed, nonparametric analyses were applied. Depending on the type of data, group comparisons were made using the chi-square test, Student’s t-test, Kruskal-Wallis test, or one-way ANOVA, where appropriate. All statistical analyses were performed using R Software version 4.1.2 (2021-11-01) for macOS. All P values were two-tailed, and a P value <0.05 was considered statistically significant.

## Results

A total of 122 individuals were included in this study, comprising 62 healthy controls (Control_A= 37; Control_B= 25), 15 adenoma cases, and 45 colorectal cancer (CRC) cases, including 23 right-sided (CRC_R) and 22 left-sided (CRC_L) tumors. Individuals in the Control_B subgroup were significantly younger and leaner than those in the other subgroups. (**Table 1**). The distribution of enterotypes differed significantly among the control, adenoma, and CRC groups (χ² = 12.97, df = 4, p = 0.011). The CRC group showed a higher proportion of the Bacteroides/Phocaeicola (62.2%) and Prevotella (26.7%) enterotypes, while the Firmicutes enterotype was relatively enriched in the adenoma (46.7%) and control groups (37.1%). The Prevotella enterotype was least prevalent in controls (11.3%) and most prevalent in the CRC group (26.7%) (**Table 2**, **Figure 1**).

**Figure 1.**
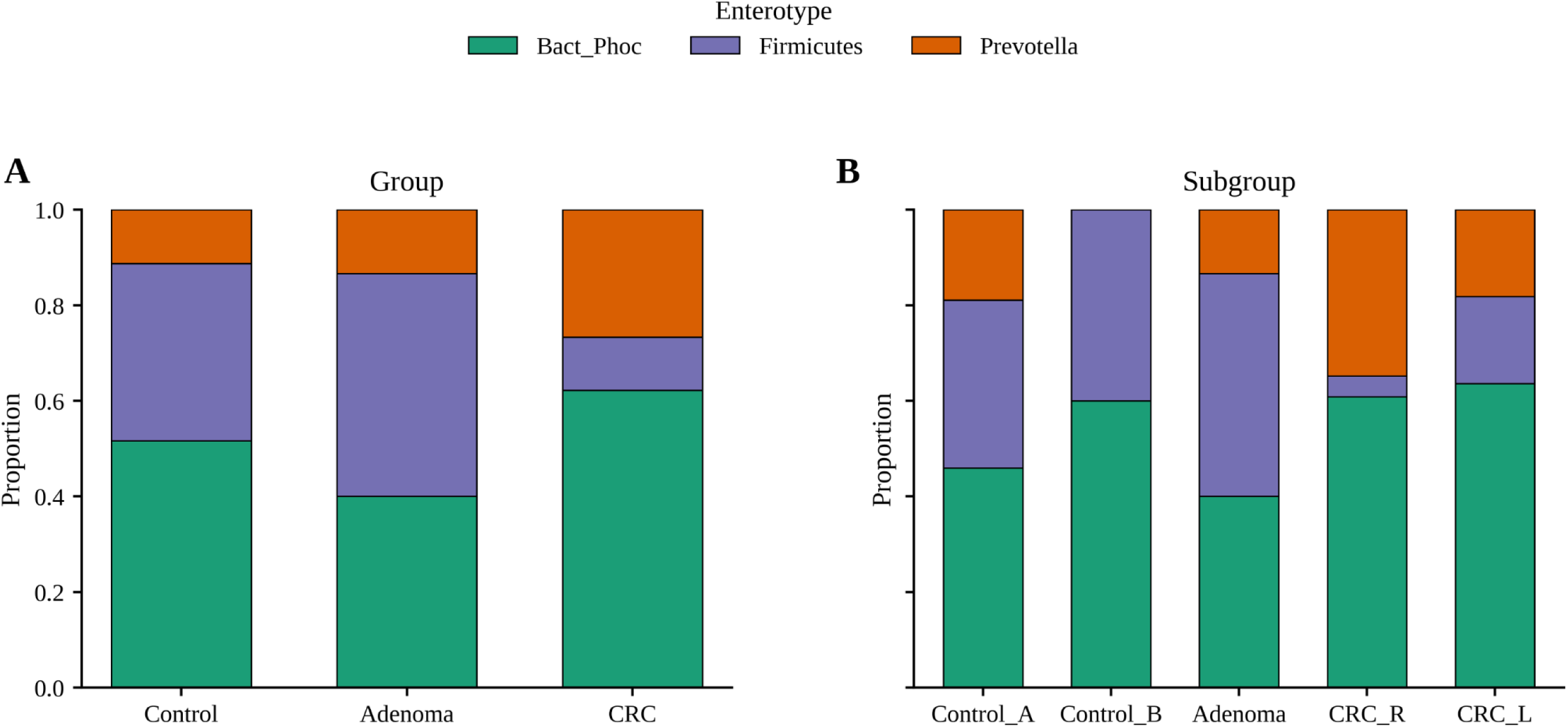
**Enterotype distribution across groups.**

**Table 1.**
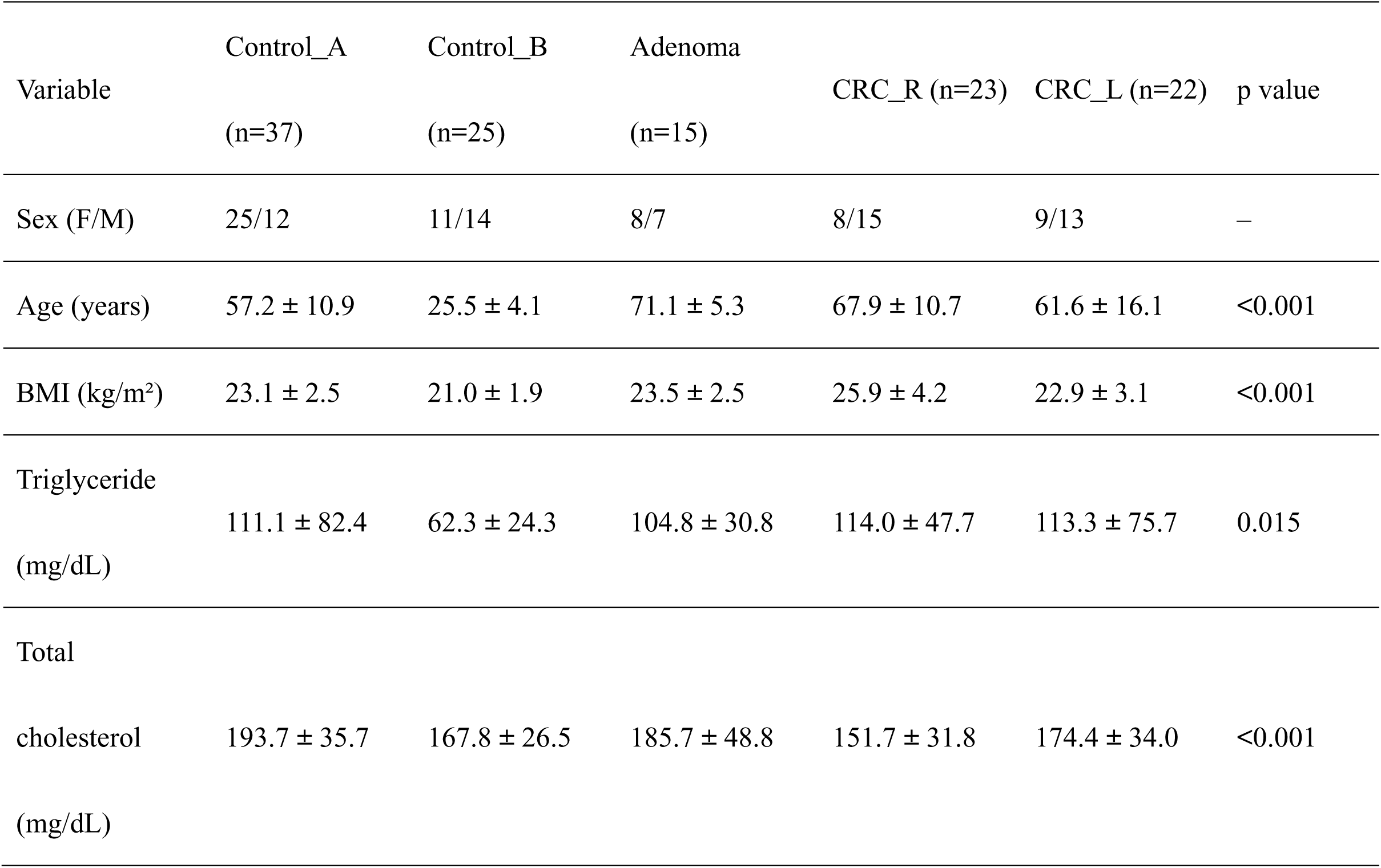
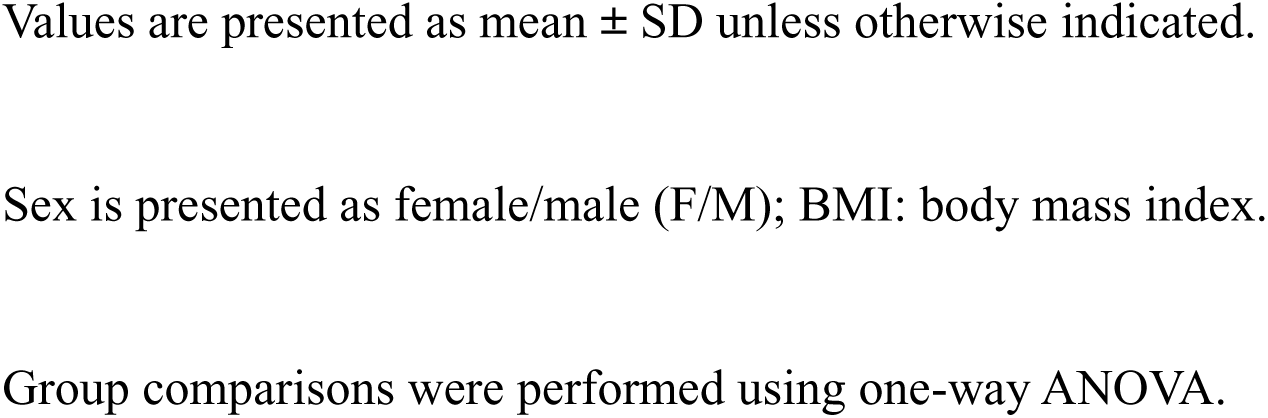
Baseline characteristics of the study subjects.

**Table 2.** Enterotype distribution across groups.

| Group | Bact_Phoc n (%) | Firmicutes n (%) | Prevotella n (%) | Total |
| --- | --- | --- | --- | --- |
| Control | 32 (51.6) | 23 (37.1) | 7 (11.3) | 62 |
| Adenoma | 6 (40.0) | 7 (46.7) | 2 (13.3) | 15 |
| CRC | 28 (62.2) | 5 (11.1) | 12 (26.7) | 45 |
| Total | 66 (54.1) | 35 (28.7) | 21 (17.2) | 122 |

8 Bact\_Phoc: Bacteroides/Phocaeicola; Total: total number or total subjects

### Alpha- and Beta-Diversity Analyses

Alpha diversity indices including observed features index, Faith PD, Pielou’s evenness, and the Shannon diversity showed no significant difference between the groups and subgroups (**Figure 2**).

**Figure 2.**
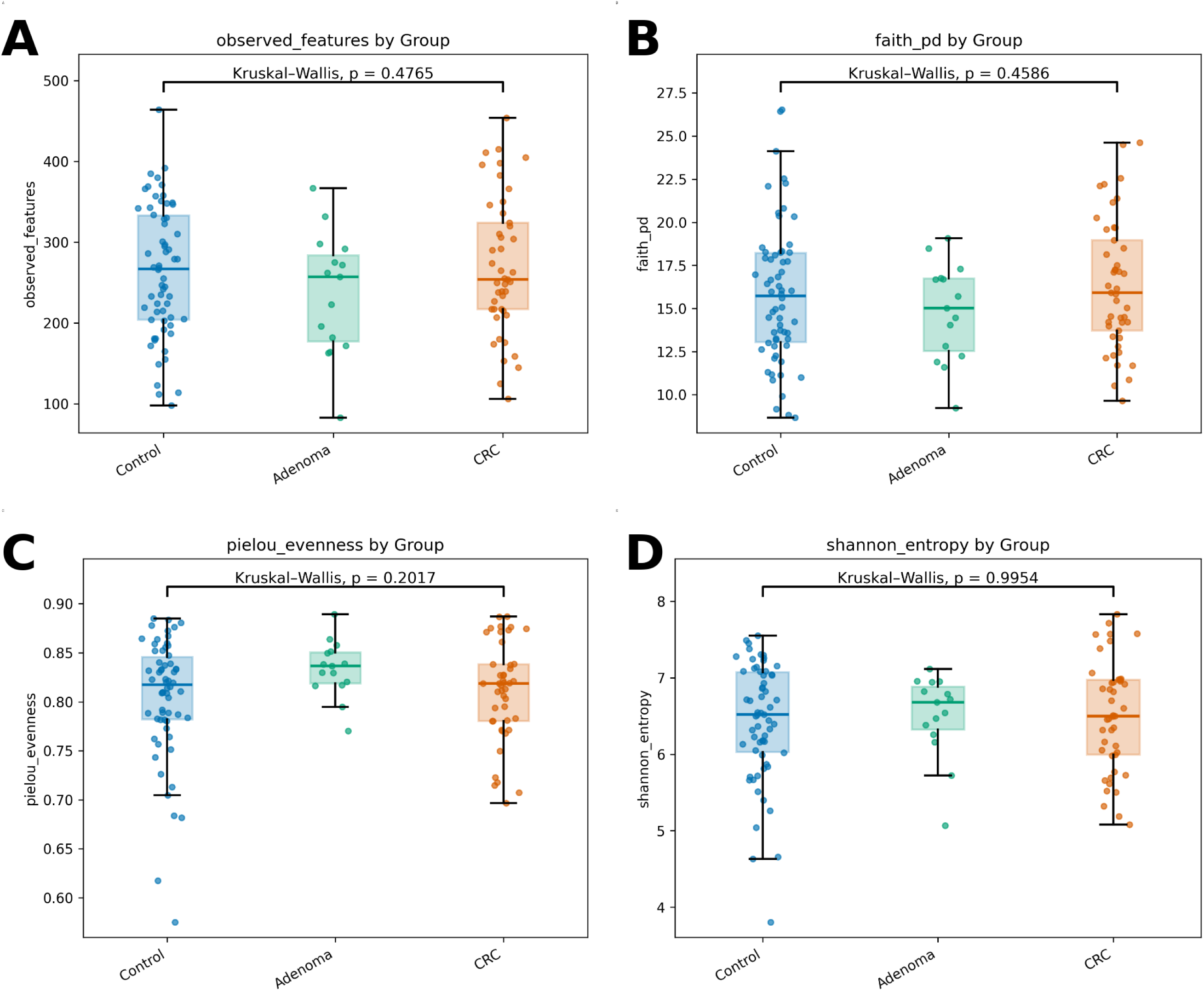
The alpha diversity of fecal microbiota between patients with colorectal cancer, adenoma and controls. Observed features, (2A), Faith’s phylogenetic diversity (Faith PD, 2B), Pielou’s evenness index (2C), and the Shannon diversity index (2D) among three groups. The boxes (containing 50% of all values) show the median (horizontal line across the middle of the box) and the interquartile range, whereas the blackspots represent the 10^th^ and the 90^th^ percentiles. CRC = colorectal cancer

Bray–Curtis-based beta diversity analysis revealed significant differences in overall gut microbial community composition across both the three major groups and the five subgroups. At the group level, PERMANOVA demonstrated a significant difference among Control, Adenoma, and CRC (R² = 0.021, F = 1.287, p < 0.001). In the subgroup analysis, the PERMANOVA result remained significant (R² = 0.041, F = 1.251, p < 0.001), indicating that subgroup classification explained 4.1% of the total variance.

Pairwise analysis at the group level showed that all comparisons were significant, including Control vs CRC (R² = 0.013, p < 0.001), Adenoma vs CRC (R² = 0.019, p = 0.006), and Control vs Adenoma (R² = 0.017, p < 0.001). However, pairwise PERMDISP was significant for both Control–Adenoma and Adenoma–CRC, suggesting that these differences were influenced, at least in part, by unequal within-group variability. In contrast, the Control–CRC comparison remained significant without a significant pairwise PERMDISP result (p = 0.055), supporting a more robust difference in community composition between these two groups.

At the subgroup level, Adenoma was not significantly different from CRC_R (R² = 0.029, p = 0.088) but remained significantly different from CRC_L (R² = 0.032, p = 0.001), indicating that adenoma may share partial microbial features with specific CRC subgroups rather than with CRC. Remarkably, Control_B showed significant differences from Adenoma, CRC_R, and CRC_L in pairwise PERMANOVA (all p < 0.001), and these differences were not accompanied by significant pairwise PERMDISP results, suggesting more robust compositional separation. By contrast, Control_A showed significant pairwise differences from all other subgroups; however, these comparisons were also accompanied by significant pairwise PERMDISP results, indicating that the observed separation may reflect both centroid differences and heterogeneity of within-group variability. PCoA plots showed partial clustering trends, although substantial overlap remained among groups (**Figure 3**).

**Figure 3.**
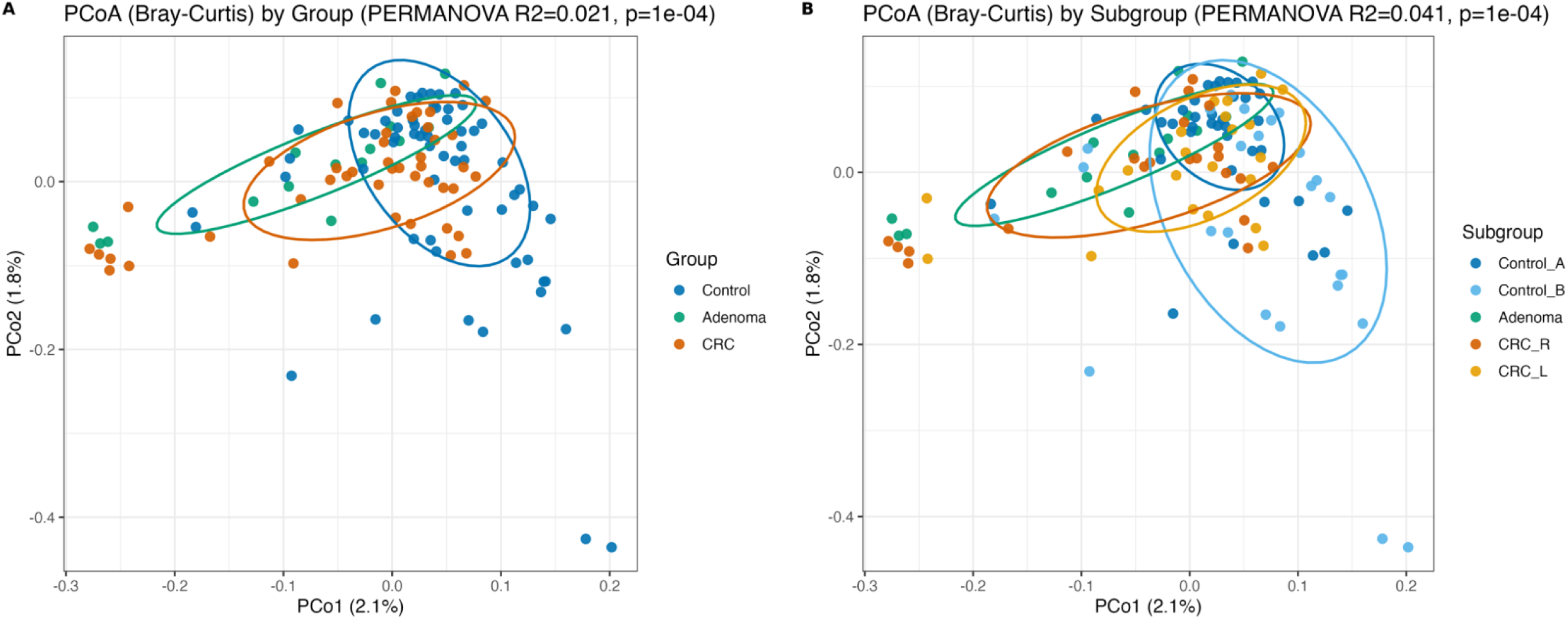
Bray-Curtis dissimilarity and principal coordinate analysis (PCoA) of bacterial abundance. CRC = colorectal cancer

### Differential Abundance Analyses

At the genus level, Blautia, Alistipes, and Anaerobutyricum were more abundant in the control group, with the highest relative abundance observed in the Control_B subgroup. In contrast, the adenoma group exhibited relatively higher abundances of Lachnospira and Agathobacter compared with the other groups. Meanwhile, the CRC group was characterized by a marked enrichment of Fusobacterium and Parabacteroides. (**Suppl. Figure 1**).

LEfSe analysis (LDA score > 2.0) identified Prevotella as significantly enriched in the CRC group and Ruminococcus in the adenoma group. At the subgroup level, *Fusicatenibacter saccharivorans* was enriched in Control_B, whereas *Phascolarctobacterium faecium* was more abundant in the adenoma subgroup. No other taxa exceeded the significance threshold (**Suppl. Figure 2**).

ANCOM-BC2 analysis identified several taxa that potentially associated with colorectal tumorigenesis. *Lawsonibacter sp900066825, Bacteroides luhongzhoulii, Ruminococcus B gnavus*, and *Phocaeicola vulgatus*, were significantly enriched in adenoma compared with controls. Several taxa exhibited progressive abundance changes along the adenoma–carcinoma sequence. *Phocaeicola dorei, Anaerotignum faecicola, Negativibacillus massiliensis*, and *Dysosmobacter segnis* revealed increasing abundance from control to adenoma and CRC, suggesting potential involvement in tumor progression. On the other hand, beneficial butyrate-producing bacteria, including *Agathobacter rectalis* and *Faecalibacillus intestinalis*, showed decreasing abundance toward CRC. (**Figure 4, 6, Suppl. Figure 3-5**). After adjustment for age, BMI, and sex, *Lawsonibacter sp900066825* remained significantly enriched in the adenoma group.

**Figure 4.**
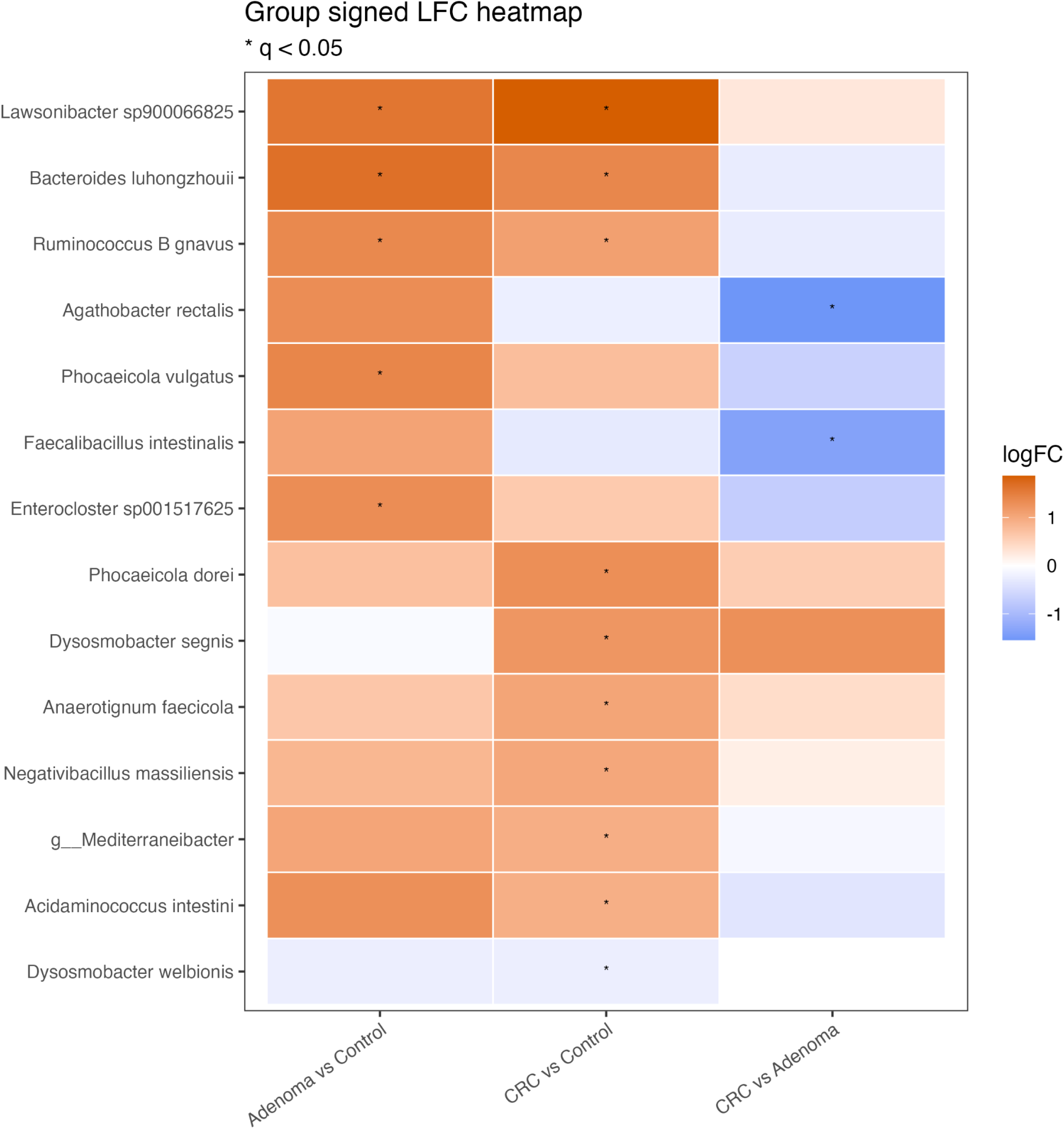
Differentially abundant bacterial taxa across the adenoma–carcinoma sequence. Heatmap showing the signed log fold change (logFC, LFC) of representative bacterial taxa identified by differential abundance analysis using ANCOM-BC2. Positive values (orange) indicate higher abundance in the first group of each comparison, whereas negative values (blue) indicate higher abundance in the second group. Pairwise comparisons include adenoma vs control, CRC vs control, and CRC vs adenoma. Asterisks indicate statistically significant differences after false discovery rate correction (q < 0.05).

### Function Prediction

Functional pathway analysis using PICRUSt2 and ANCOM-BC2 revealed significant metabolic alterations across the adenoma–carcinoma sequence, consistent with the progressive microbial shifts observed at the taxonomic level. Compared with controls, the adenoma group showed enrichment of several metabolic pathways, particularly those related to amino acid metabolism, including the threonine catabolism pathway (THREOCAT-PWY) and the methylglyoxal degradation pathway (METHGLYUT-PWY), as well as fermentation-related pathways, such as PWY-5181, PWY-6185, and PWY-5417. In addition, pathways involved in aromatic compound degradation, including the catechol ortho-cleavage pathway (CATECHOL-ORTHO-CLEAVAGE-PWY) and the protocatechuate ortho-cleavage pathway (PROTOCATECHUATE-ORTHO-CLEAVAGE-PWY), were also enriched in adenoma (**Figure 5, 6, Suppl. Figure 3-5**).

**Figure 5.**
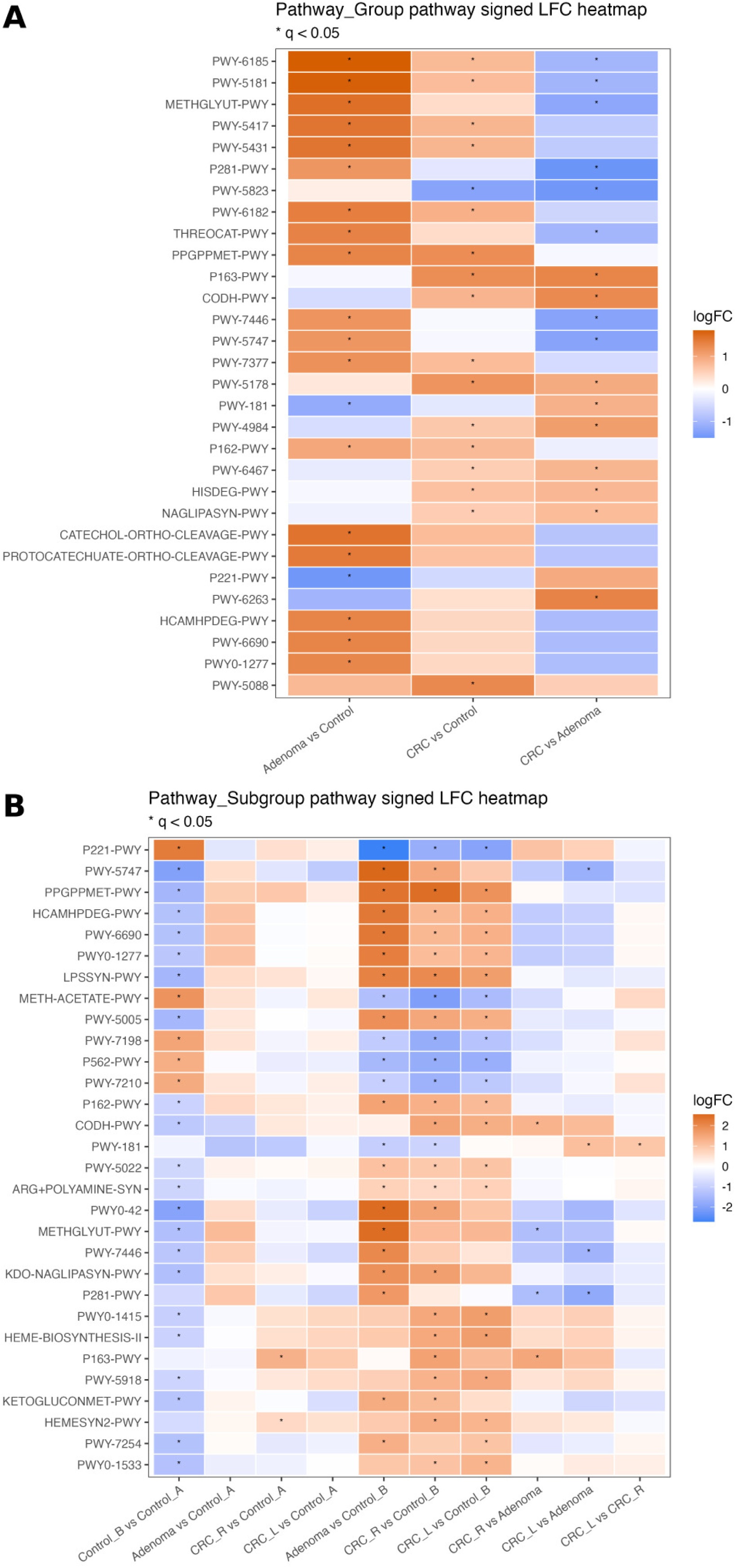
Differentially abundant microbial pathways across groups and subgroups. (A) Heatmap showing the signed log fold change (logFC, LFC) of microbial metabolic pathways across pairwise comparisons among the control, adenoma, and colorectal cancer (CRC) groups. (B) Heatmap showing pathway differences across clinical subgroups, including Control_A, Control_B, Adenoma, CRC_R, and CRC_L. Functional pathway abundance was inferred using PICRUSt2 and differential analysis was performed using ANCOM-BC2. Positive values (orange) indicate higher pathway abundance in the first group of each comparison, whereas negative values (blue) indicate higher abundance in the second group. Asterisks indicate statistically significant differences after false discovery rate correction (q < 0.05).

**Figure 6.**
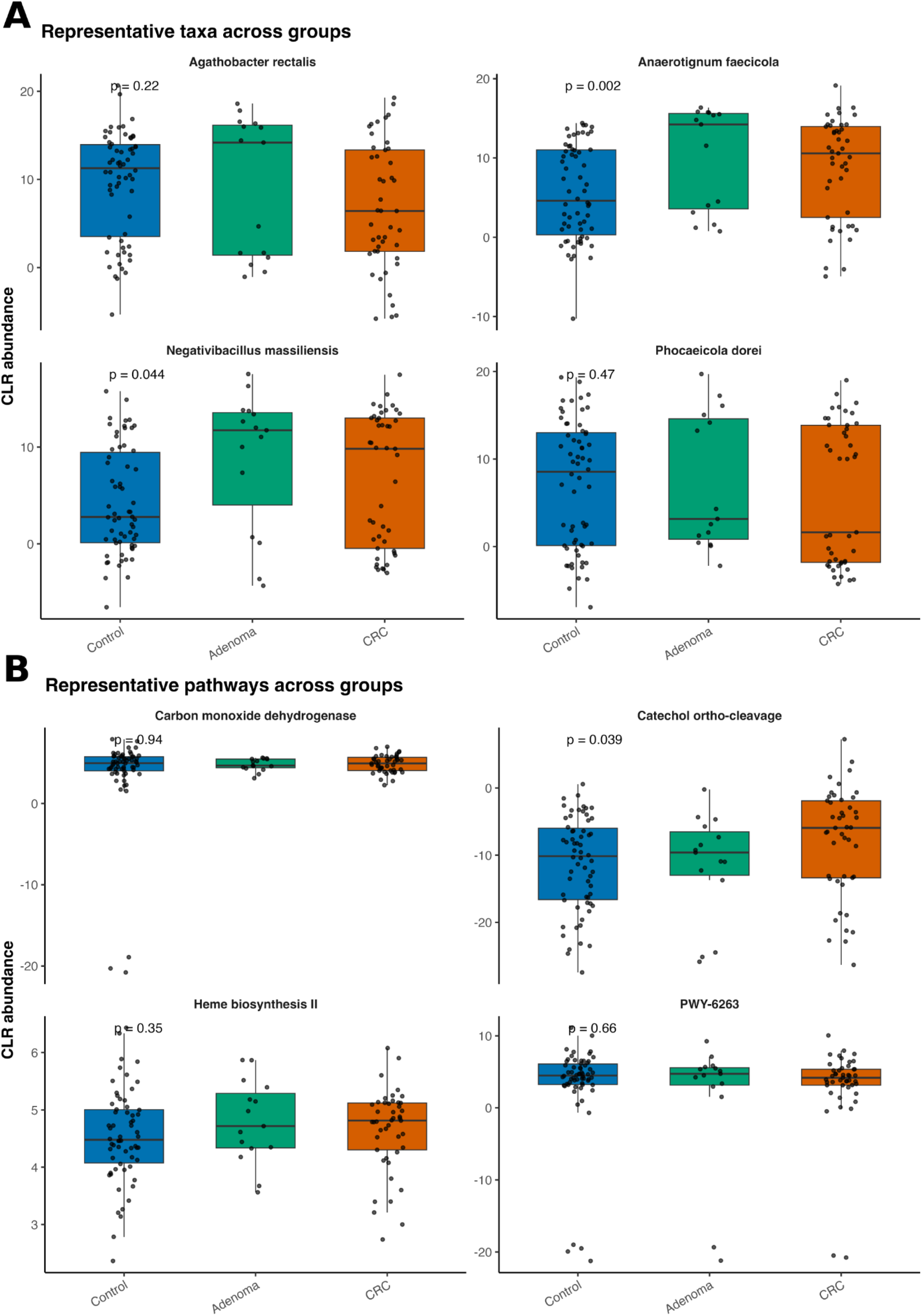
Representative microbial taxa and metabolic pathways across the adenoma–carcinoma sequence. (A) Centered log ratio (CLR)-transformed abundance of representative bacterial taxa across control, adenoma, and colorectal cancer (CRC) groups. (B) CLR-transformed abundance of representative microbial pathways predicted by PICRUSt2 across groups. Overall p-values were calculated using the Kruskal–Wallis test. Boxes indicate interquartile ranges with medians, and dots represent individual samples.

In contrast, the CRC group exhibited enrichment of pathways associated with energy metabolism and bacterial stress responses, including the carbon monoxide dehydrogenase pathway (CODH-PWY), PWY-6263, P163-PWY, and the heme biosynthesis II pathway (HEME-BIOSYNTHESIS-II). These findings suggest a transition from metabolically diverse microbial activity in adenoma to a more specialized metabolic profile in CRC. Notably, several pathways enriched in both adenoma and CRC, including PWY-5181, PWY-6185, PWY-5417, the catechol ortho-cleavage pathway, protocatechuate ortho-cleavage pathway, CODH-PWY, and PWY-6263, showed positive correlations with Klebsiella, whereas the heme biosynthesis II pathway was positively associated with Prevotella (Figure 5-7, Suppl. Figure 3-5).

**Figure 7.**
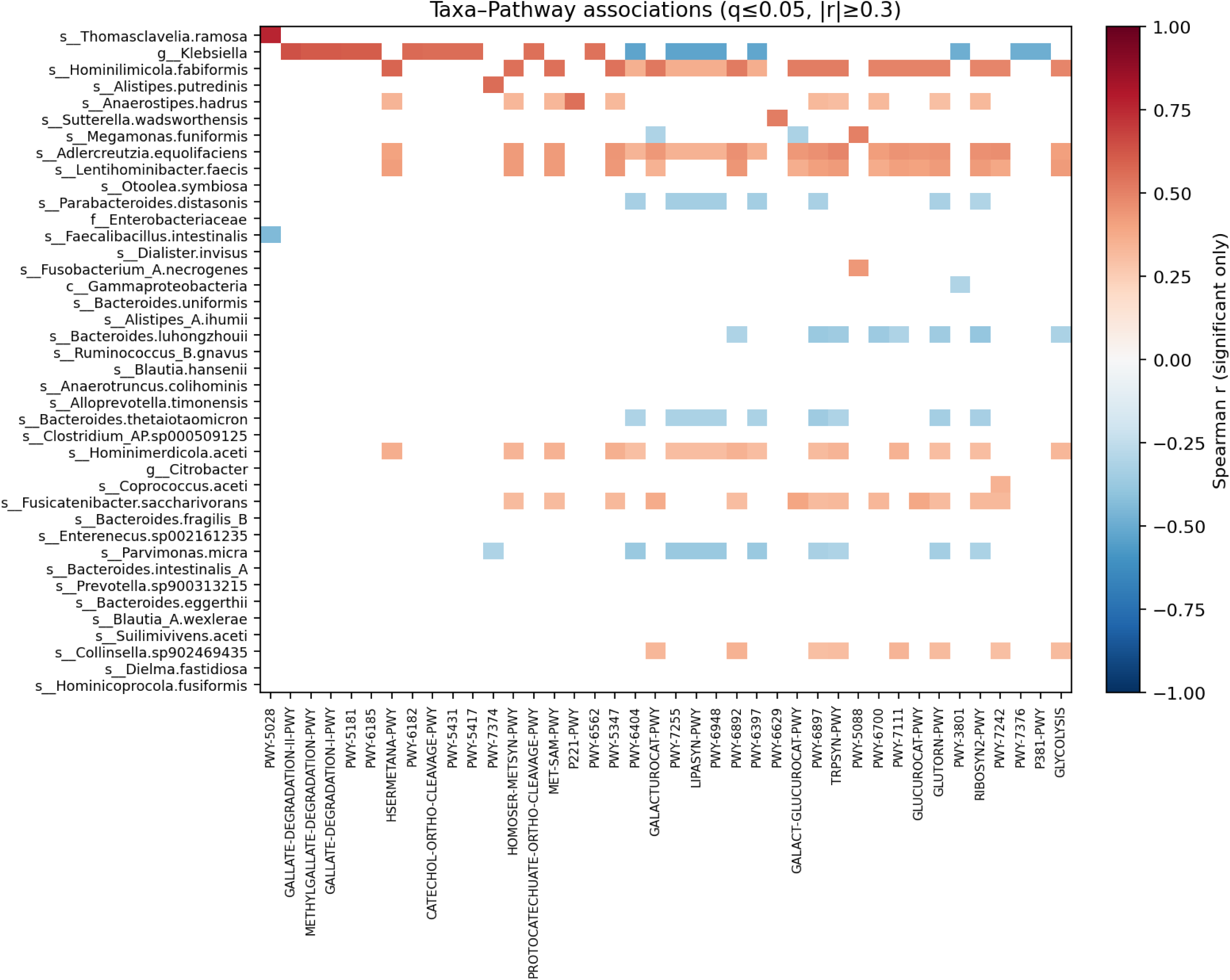
Taxa–pathway correlation network across the gut microbiome. Heatmap illustrating significant associations between bacterial taxa and predicted microbial metabolic pathways. Spearman’s rank correlation analysis was used to evaluate relationships between taxa and pathways. Only correlations meeting the significance thresholds (q ≤ 0.05 and |r| ≥ 0.3) are shown. Positive correlations are indicated in red and negative correlations in blue, with color intensity corresponding to the magnitude of the correlation coefficient. Taxonomic annotations are shown at the species level when available, and functional pathways were predicted using PICRUSt2.

Subgroup analysis further revealed functional heterogeneity within the control cohort. Several pathways related to amino acid metabolism and microbial stress responses, including PWY-5747, the ppGpp biosynthesis pathway (PPGPPMET-PWY), the homocysteine and cysteine metabolism pathway (HCAMHPDEG-PWY), PWY-6690, PWY0-1277, the methylglyoxal degradation pathway (METHGLYUT-PWY), and PWY-7446, were more abundant in the Control_A subgroup compared with Control_B. These pathways showed positive correlations with multiple bacterial taxa, including members of Gammaproteobacteria and genera such as Succinivibrio, Alloprevotella, Caproiciproducens, and Dorea. This functional heterogeneity may partly reflect host-related factors, as individuals in the Control_B subgroup were generally younger and leaner, whereas the Control_A subgroup represented a more heterogeneous population (**Figure 5, 6**).

In addition, several pathways involved in microbial stress response and amino acid metabolism (PPGPPMET-PWY, METHGLYUT-PWY, and HCAMHPDEG-PWY) were significantly increased in the adenoma group compared with both control subgroups, suggesting enhanced microbial metabolic activity during early tumorigenesis.

### Metabolomic analysis

A total of 61 stool samples were subjected to metabolomic analysis, including 21 from Control_A, 15 from the adenoma group, 11 from CRC_R, and 14 from CRC_L. Fecal short-chain fatty acid (SCFA) profiles differed significantly across groups. Formic acid decreased from 182.4 ± 150.8 in controls to 19.2 ± 55.1 in the CRC group, whereas the adenoma group showed higher levels (360.5 ± 213.1). Similar trends were observed for other major SCFAs, including acetic acid (2190.6 ± 1680.8 in controls vs 699.0 ± 796.7 in CRC vs 1263.6 ± 924.5 in adenoma), propionic acid (742.7 ± 656.1 vs 180.1 ± 230.6 vs 327.4 ± 289.3), and butyric acid (205.0 ± 153.1 vs 11.6 ± 13.7 vs 74.8 ± 66.3). Formic acid remained similarly low in both left- and right-sided CRC (22.8 ± 19.1 and 15.7 ± 7.4, respectively) compared with controls (182.4 ± 187.0). In the subgroup analysis, acetate, propionate, butyrate, and formate were significantly higher in the adenoma group than in controls and both CRC subgroups (Kruskal–Wallis FDR q ≤ 2.8×10⁻⁴; post-hoc BH-adjusted q ≤ 0.038). Within CRC, CRC_R showed higher acetate, propionate, and butyrate than CRC_L (pairwise q = 0.013, 0.049, and 0.011, respectively).

Bile acid profiling identified tauro-ursocholanic acid as the most discriminative metabolite, with markedly higher levels in both CRC subgroups compared with controls and adenoma (Kruskal–Wallis FDR q = 4.3×10⁻⁸; post-hoc: CRC_R vs control q = 3.9×10⁻⁶, CRC_L vs control q = 0.0039, CRC_R vs adenoma q = 2.3×10⁻⁵, CRC_L vs adenoma q = 1.2×10⁻⁴). In contrast, 3α-hydroxy-6,7-diketo cholanic acid and lithocholic acid were reduced in adenoma relative to controls and CRC (pairwise q = 0.0048–0.036 and q = 0.022–0.047, respectively). Other bile acids showed substantial inter-individual variability (e.g., 3-ketocholanic acid/dehydrolithocholic acid) and did not demonstrate robust between-group differences after multiple-testing correction.

To characterize coordinated microbiome–metabolite structure, we performed sPLS analysis on residualized genus-level CLR abundances and metabolite panel features after adjustment for group, age, BMI, and sex. The model identified a dominant bile acid–related latent axis, with total bile acids (BA_total) and unconjugated bile acids (BA_unconjugated) showing highly concordant directions. *Faecalimonas* and *Enterocloster* were aligned with this bile acid axis, whereas *Prevotella* and *Faecalibacterium* showed discordant or opposite orientations (**Suppl. Figure 6A**). In the sample space, the control, adenoma, and CRC groups showed partial overlap, indicating that the integrated signature primarily captured shared covariance between taxa and metabolites rather than strong group discrimination (**Suppl. Figure 6B**). These multivariate findings were concordant with the pairwise partial correlation analysis, in which *Faecalimonas* was positively associated with BA_total and BA_unconjugated (rho = 0.355, q = 0.0956 for both), while *Alitiscatomonas* was positively associated with total SCFAs (Total_SCFA) (rho = 0.346, q = 0.0956) (**Suppl. Figure 7**).

## Discussion

In this study, we evaluated gut microbial composition and predicted functional profiles in patients with colonic adenoma and CRC using full-length 16S rRNA sequencing. Although overall community diversity and microbial structure showed only modest differences across groups, several specific taxa and predicted metabolic pathways exhibited significant differential distributions. Furthermore, both taxonomic and functional alterations were observed along the adenoma–carcinoma sequence, suggesting progressive microbial shifts during colorectal tumorigenesis. In addition, despite the relatively limited sample size for metabolomic profiling, distinct metabolic patterns were identified: tauro-ursocholanic acid was enriched in CRC samples, whereas some SCFAs, including acetic acid, propionic acid, and butyric acid, showed a stepwise decrease from controls to adenoma and CRC.

In this study, several microbial taxa demonstrated progressive abundance changes across the control, adenoma, and CRC groups, suggesting their potential involvement in colorectal tumorigenesis and disease progression. Notably, *Phocaeicola dorei*, *Anaerotignum faecicola*, *Negativibacillus massiliensis*, and *Dysosmobacter segnis* exhibited increasing abundance from controls to adenoma and CRC, indicating a potential association with disease progression. Previous studies have reported enrichment of *Phocaeicola dorei* and *Anaerotignum faecicola* in CRC-associated microbiomes, ^5,27–29^ whereas *Negativibacillus massiliensis* has been proposed as a marker of healthy microbial communities in a machine learning–based CRC classification model, suggesting that certain microbial taxa may play context-dependent roles in colorectal carcinogenesis. ^30^ Furthermore, *Faecalibacillus intestinalis*, which has been reported to be reduced in several types of cancer, ^28^ showed higher abundance in the adenoma group than in the CRC group in the present study, suggesting a potential decline during the adenoma–carcinoma transition. In contrast, *Lawsonibacter* sp900066825, *Bacteroides luhongzhoulii*, *Ruminococcus B gnavus*, and *Phocaeicola vulgatus* were enriched in both the adenoma and CRC groups compared with controls, although these taxa did not exhibit a clear progressive trend across disease stages.

Despite the microbial shifts observed along the adenoma–carcinoma sequence, corresponding alterations in microbial metabolic functions were also expected. Due to the limitations of amplicon sequencing, PICRUSt2 was used to predict functional profiles of the gut microbiome. Several pathways enriched in the adenoma group were primarily associated with amino acid metabolism and microbial fermentation processes, including threonine catabolism and methylglyoxal degradation pathways.^25,26,31^ In addition, pathways involved in aromatic compound degradation, such as the catechol and protocatechuate ortho-cleavage pathways, were also increased in the adenoma group, suggesting active microbial metabolic adaptation during early tumorigenesis. In contrast, the CRC group exhibited enrichment of pathways related to energy metabolism and bacterial stress responses, including carbon monoxide dehydrogenase pathways and heme biosynthesis. ^25,26,31^ These functional alterations may reflect adaptive metabolic responses of the gut microbiome to the evolving tumor-associated intestinal microenvironment during colorectal tumorigenesis. Consistent with these findings, metabolomic analysis revealed that the major SCFAs gradually decreased from controls to adenoma and CRC, whereas tauro-ursocholanic acid was elevated in CRC samples, indicating a shift in microbial metabolic outputs during colorectal tumorigenesis.

Furthermore, two distinct healthy control cohorts were included in this study, and considerable heterogeneity in age and metabolic profiles was observed between them. Pathways associated with amino acid metabolism and microbial stress responses, including the ppGpp biosynthesis pathway, homocysteine and cysteine metabolism pathway, and methylglyoxal degradation pathway, were more abundant in the Control_A subgroup and further increased in the adenoma group. This progressive pattern suggests that individuals in Control_A may represent an intermediate metabolic state between healthy and pre-disease conditions. These findings also highlight that host metabolic status may influence gut microbial functional profiles even within presumably healthy populations.

Several limitations of this study should be acknowledged. First, as this study employed a cross-sectional design, establishing temporal changes in gut microbiota along the adenoma–carcinoma sequence remains challenging. Consequently, the relationship between microbial alterations and disease progression, severity, or prognosis could not be determined. Second, detailed information on potential confounding factors, such as habitual diet and lifestyle, was not systematically documented, which may have influenced the interpretation of the microbiome findings. Third, the control cohort exhibited heterogeneity in metabolic profiles, which may have partially confounded the observed microbial shifts from controls to adenoma and CRC. In addition, functional profiles were inferred using PICRUSt2 based on 16S rRNA gene sequencing data, which may not fully capture the true functional capacity of the gut microbiome. Future studies incorporating shotgun metagenomics and metabolomic analyses in larger and more metabolically homogeneous cohorts will be necessary to further elucidate the mechanistic role of gut microbial metabolism in colorectal tumorigenesis.

In conclusion, this study identified taxonomic and functional shifts in the gut microbiome along the adenoma–carcinoma sequence using full-length 16S rRNA sequencing. Functional prediction suggested increased microbial activity related to amino acid metabolism, aromatic compound degradation, and stress-response pathways during colorectal tumorigenesis. These findings were further supported by metabolomic analysis, which demonstrated a gradual decrease in short-chain fatty acids and an increase in bile acid–related metabolites in CRC. These results suggest that alterations in gut microbial composition and metabolism may contribute to colorectal tumor development and provide potential insights for microbiome-based biomarkers and preventive strategies.

## Author Contribution

TEC and JCL wrote the main manuscript text. TEC performed the analysis and prepared all the figures. TEC, HHL, JCL, KCL, PCL, YTL, YFC and YPW helped to enrolled participants. JCL, HCH, CWS, YHH and MCH designed the research. JCL provided the funding acquisition. All authors reviewed the manuscript.

